# Fitness effects of killer virus infection on wild *Saccharomyces paradoxus*

**DOI:** 10.1101/2024.09.06.611503

**Authors:** Rahul Unni, Onur Erk Kavlak, Eva H. Stukenbrock, Primrose J. Boynton

## Abstract

Endosymbioses, the intimate relationships between smaller symbionts and larger hosts, have profound impacts on eukaryotic organisms. However, symbiont effects on host fitness in natural conditions are difficult to study, especially for microbial hosts. We used killer viruses and the wild yeast *Saccharomyces paradoxus* to study a symbiotic virus’s effect on its host’s fitness in oak litter. We cured hosts of naturally-occurring killer viruses and compared killer and cured individuals’ fitnesses in laboratory medium and oak litter using a unique field chamber design to house competing *S. paradoxus*. In the laboratory, the impact of virus loss on host fitness could be positive, negative, or neutral, depending on host identity. Trends in the forest were similar to those in the lab, although only overall strain fitness differences were significant and curing impacts differed between the forest and laboratory. These results demonstrate the importance of incorporating environmental context into studies of host-symbiont interactions.

## Introduction

Endosymbioses are ubiquitous features of life. These intimate relationships involve one or more smaller partners (symbionts) living within a larger partner (the host) (Bright & Bulgheresi, 2010). Hosts range in size from viruses hosting virophages, through bacterial cells hosting plasmids and phages, all the way up to plants and animals hosting diverse microbiomes (Castledine & Buckling, 2024; del Arco et al., 2024; Hassani et al., 2019; Lloyd & Thomas, 2023; Lozupone et al., 2012). Endosymbiosis impacts are substantial and include the radiation of all eukaryotic organisms after the evolution of endosymbiotic mitochondria (Gabaldón, 2021). Endosymbionts have profound impacts on host fitness, both positive and negative, and these impacts depend on host or symbiont identities or environmental conditions. For example, curing *Drosophila melanogaster* of infecting Wolbachia endosymbionts has different effects on host survival and fecundity depending on host genotype (Fry et al., 2004). Additionally, arbuscular mycorrhizal fungi can act as parasites in high-phosphorus conditions and mutualists in low-phosphorus conditions (Johnson, 1998). The relationships among environment, endosymbiosis, and fitness are easiest to establish with macrobial hosts in controlled laboratory experiments. These relationships are much more difficult to establish in microbial hosts in natural environments such as forests or grasslands, in part because microbes are difficult to track *in situ*. Instead, most investigations of microbial host fitness are conducted in laboratories (*e*.*g*., bacterial-plasmid and bacterial-phage interactions) (Obeng et al., 2016; San Millan & MacLean, 2017; Shapiro & Turner, 2018). In this study, we used field chambers to understand the fitness consequences of endosymbiosis in a tractable model organism in a forest soil environment.

The *Saccharomyces* killer system, in which double-stranded RNA (dsRNA) viruses live inside single-celled yeast hosts, has long been a model of endosymbiosis (Rowley, 2017). Members of the genus *Saccharomyces*, which includes the famous laboratory model *S. cerevisiae*, can have a diversity of cytoplasmic viruses; one of these, a “killer virus,” produces transcripts that code for secreted toxins that can kill nearby sensitive yeasts (Wickner, 1996). Killer viruses depend on helper viruses, which coexist with killer viruses in the host cytoplasm, for their maintenance and replication (Schmitt & Breinig, 2006). The viruses are transmitted through cell division and mating and cannot enter cells through the cell wall. Killer toxins are widespread among single-celled fungi (*i*.*e*., yeasts broadly defined) in general, and may play important ecological roles by killing competing yeasts (Boynton, 2019; Crabtree et al., 2023). These toxins are coded on a variety of genetic elements in different yeast species, including cytoplasmic viruses, plasmids, and the cell’s nuclear genome, although dsRNA viruses in *Saccharomyces* are probably the best studied (Boynton, 2019; Rowley, 2017).

We do not yet understand the fitness consequences of the killer virus endosymbiosis, especially in non-domesticated *Saccharomyces* habitats. In laboratory experiments, toxin production appears to have intrinsic fitness costs (Pintar & Starmer, 2003). These fitness costs can be countered through host-symbiont coevolution and extrinsic benefits of toxin production in a community context, such as competitor interference (Boynton, 2019; Pieczynska et al., 2017; Wloch-Salamon et al., 2008). However, the costs and benefits of killer virus endosymbiosis have not yet been explored in natural environments. For example, while members of the genus *Saccharomyces* are laboratory models and are common in human fermentation environments, they grow wild and most likely evolved in forest environments (Boynton & Greig, 2014). A much larger *Saccharomyces* diversity exists on forest substrates, such as soil, leaf litter, and tree bark, than exists in laboratories or fermentations (Boynton & Greig, 2014; Wang et al., 2012), and the forest environment is different (e.g., more heterogeneous; more carbon limited; often involving a greater diversity of interacting organisms) from human environments (El-Khatib et al., 2023). The costs and benefits of hosting killer endosymbionts may therefore vary considerably between the laboratory and the forest.

Here we used the wild yeast *Saccharomyces paradoxus* to investigate the intrinsic impacts of killer viruses on host fitness in laboratory and natural environments. We isolated killer, sensitive, and resistant *S. paradoxus* from soil and leaf litter in a temperate forest as part of a previous study (Boynton et al., 2021). We compared relative fitness (relative to a toxin-resistant control *S. paradoxus*) of killer yeasts with and without their endosymbiotic viruses. By using a toxin-resistant control as a reference, we removed impacts of the secreted toxin from fitness assays. We asked: 1) whether virus curing increased or decreased host fitness; 2) if fitness effects of curing were consistent across *S. paradoxus* isolates; and 3) if the fitness effects of curing were consistent in laboratory and field environments. We found that the effect of curing on host fitness varied depending on both environment and host identity.

## Methods

### *S. paradoxus* killer and cured strains used in this study

We measured fitnesses of killer *S. paradoxus* strains, killer strains that we had cured of the killer virus, and naturally-occurring nonkiller *S. paradoxus* strains from a collection of wild strains previously isolated from oak-associated litter and soil in the Nehmten forest in Schleswig-Holstein, Germany (Boynton et al., 2021) (Supplemental Table 1). Our toxin-resistant control strains were taken from the same collection. We chose fourteen killer strains from this collection; these strains had previously been confirmed to produce killer toxins using a phenotypic “halo assay” (Boynton et al., 2021). Briefly, the authors pipetted small volumes of liquid cultures of each *S. paradoxus* isolate onto lawns of sensitive *S. cerevisiae* cells and observed inhibition of sensitive lawn growth. We cured each selected killer strain of its killer virus by treatment with cycloheximide (Fink & Styles, 1972). Strains were grown to stationary phase in liquid YEPD (10 g/L yeast extract, 20 g/L peptone, and 20 g/L dextrose) and then diluted with sterile water and plated onto solid YEPD medium (YEPD with 25 g/L agar) with 13.3 µg/plate of cycloheximide. The resulting colonies were then tested for killing by replica plating onto sensitive *S. cerevisiae* lawns and looking for inhibition of lawn growth. Colonies that showed no killing activity were identified as cured strains. Colonies from the same petri dish that survived the cycloheximide curing treatment but continued to show killing activity were chosen for the fitness assays to control for the effects of the curing process. Cycloheximide treatment generally cures killer yeasts of the killer virus but not the helper virus (Ball et al., 1984). A list of cured and control *S. paradoxus* strains is provided in supplementary table 1.

### Laboratory fitness assay

To understand how curing the virus affects host fitness in a laboratory environment, we first performed a laboratory fitness assay with eight killer strains and their corresponding cured versions. We measured the reproductive fitness of each killer and cured isolate relative to a standard toxin-resistant reference *S. paradoxus* strain from the same environment (Lenski et al., 1991). The reference strain is resistant to all killers at 23 °C, and does not show killing activity against any tested strain. We transformed the reference strain by replacing the *mf-α2* pheromone gene with a kanamycin resistance gene using homologous recombination and standard yeast transformation procedures (Gietz & Woods, 2002). We conducted six replicate fitness assays for each of eight killer and eight cured *S. paradoxus* strains. For each fitness assay, we first conditioned the yeast isolates to the assay environment by mixing stationary phase liquid cultures of the tested and reference strains together, diluting the mixture 1:100 in liquid YEPD medium, and incubating the mixture at 15 °C for eight days. We then measured reproductive fitness by diluting the conditioned culture 1:100 again in liquid YEPD medium, counting colony-forming units of the mixture with and without kanamycin, letting the culture grow again for eight days at 15 °C, and counting colony-forming units again. Plates with fewer than 30 colonies were excluded from further analyses. We defined fitness as the natural logarithm of the ratio of numbers of tested and reference doublings (ln(tested *S. paradoxus* doublings/reference *S. paradoxus* doublings)) after eight days (Lenski et al., 1991).

### Wild fitness assay

To understand virus effects on host fitness in the host’s natural environment, we compared fitnesses of killer to corresponding cured isolates and of killer to naturally-occurring nonkiller isolates in the forest. As with the laboratory fitness assays, we tested each strain in competition with a toxin-resistant naturally-occurring reference *S. paradoxus* strain, using the change in the relative frequency of each tested strain relative to the reference strain as a proxy for reproductive fitness (Boynton et al., 2017). Instead of using a selectable marker to measure the relative growth of tested and reference strains, we tracked relative frequencies of naturally-occurring alleles at the *hxt3* locus (Boynton et al., 2017), which codes for a hexose transporter. The *S. paradoxus* population in the Nehmten forest has two alleles at this locus, and we previously used it to track changes in *S. paradoxus* frequencies in field microcosms (Boynton et al., 2017). We chose two resistant reference *S. paradoxus* strains, one of each *hxt3* genotype (*hxt3-1* and *hxt3-2*), and paired each tested strain with a reference strain of the opposite genotype. We tested twelve naturally-occurring killer strains, their twelve corresponding cured strains, and ten naturally-occurring non-killer strains (Supplementary table 1).

We conducted forest fitness assays in 12-ml tube chambers. We designed chambers (Figure 1) to ensure that the *S. paradoxus* strains studied would be in their natural environment, their environments would remain otherwise sterile, and they would not escape into the environment. Chambers were an iteration of field chambers previously used for fitness assays in the same forest (Boynton et al., 2017). For each chamber, the cap of a 12 ml polypropylene test tube (Techno Plastic Products AG, Trasadingen, Switzerland) was sliced off using a hot flame-sterilised scalpel and a sterile 25 mm diameter 0.2-µm membrane (Sartorius GmbH, Goettingen, Germany) was fixed between the cap and the test tube. The tester and reference strains were inoculated onto the membrane (2 × 10^5^ cells of each strain in 4 µl of sterile water) so that the side with the inoculated cells faced inside the test tube. The gap between the test tube and its cap was sealed using parafilm (Figure 1). We tested a total of 34 *S. paradoxus* strains (Supplementary table 1): twelve strains each of killer and corresponding cured strains, and ten nonkiller *S. paradoxus* strains from the forest. We prepared six chambers for each tested yeast strain: three replicates each for collection at two time points. Chambers were placed membrane-downward in 15-cm holes dug at the base of one of two oak trees in the Nehmten forest on the third of July 2019. Holes were then covered with soil and leaf litter. We immediately collected three replicate chambers for each strain to measure starting baseline yeast frequencies, and returned fifteen days later to collect the remaining chambers and measure final yeast frequencies.

**Figure 1:**
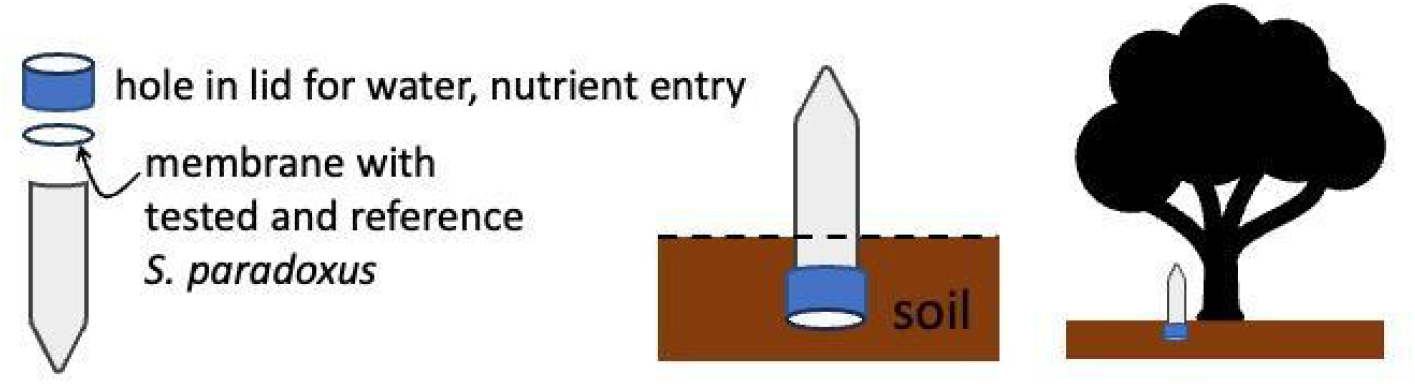
Field fitness chamber design. The top portions of test tube caps were cut off (left). Membranes with inoculated S. paradoxus strains were then clamped to the test tube mouth with the altered cap such that the inoculated strains faced inside the test tube and remained sterile (right). The opposite side of the membrane was exposed to the environment. Thus, the inoculated strains could obtain nutrients from the environment, but were not contaminated by the environment nor allowed to escape to the environment.

We extracted yeast DNA from each chamber’s membrane and then quantified tested and reference strain frequencies using digital-droplet PCR (Boynton et al., 2017). We first removed yeast cells from each membrane by vortexing it in 200 µl buffer (2% Triton X-100, 1% SDS, 100 mM NaCl, 10 mM Tris, 1 mM EDTA). Cell lysis, nucleic acid extraction with chloroform:phenol:isoamyl alcohol, and ethanol extraction of DNA were performed as previously described (Boynton et al., 2017). We then measured frequencies of the two *hxt3* alleles as previously described (Boynton et al., 2017), using dual-labelled probes and a Bio-Rad QX100 digital droplet PCR droplet generator and droplet reader (Bio-Rad, Hercules, California, USA), according to the manufacturer’s instructions. Data from droplets in which the concentration of at least one probe was less than 0.4 copies/µl were excluded. Droplet data were analysed and converted into allele frequencies using Quantasoft v. 1.7 software (Bio-Rad, Hercules, California, USA).

### Statistical analyses

We compared fitnesses among different *S. paradoxus* phenotypes (killer, cured, and non-killer) and strains using mixed-effects linear models. The dependent variable for lab assays was fitness relative to the reference strain and the dependent variable for forest assays was change in frequency of tested strain relative to the reference strain after fifteen days. Models included either experimental batch (in the lab assays) or reference strain (in the forest assays in which the reference strain needed to have a different *hxt-3* genotype from the tested strain) as a random effect. We created three models: to compare killer and cured *S. paradoxus*, we modelled the fixed effects of phenotype (killer versus cured across all strains), strain identity, and the interaction between phenotype and strain identity (differences in effects of curing across strains) in both the lab and forest; to compare killer and nonkiller S. paradoxus in the forest, we modelled the fixed effect of phenotype only. All modelling and statistical analyses were performed using R (v 3.4.3) (R Core Team, 2017) packages nlme (Pinheiro et al., 2018), lme4 (Bates et al., 2015), and lmerTest (Kuznetsova et al., 2017). Graphs were plotted using R (v 3.4.3) package ggplot2 (Wickham, 2016).

## Results

Curing neither increased nor decreased *S. paradoxus* host fitness globally; instead, the consequences of virus removal depended on assay environment and host identity. In the laboratory medium, each *S. paradoxus* strain responded to curing differently (Figure 2). Individual strains had significantly different overall fitnesses (strain: F = 4.6, df = 7,78, p = 0.0002) and changes in fitness with curing (interaction between phenotype and strain: F = 4.5, df = 7,78, p=0.0003), but overall average fitness difference between killer and cured strains was not significantly different (phenotype: F = 0.04, df = 1,78, p = 0.84). Impact of curing on fitness (*i*.*e*., differences between fitness of a host when it did not have the virus minus fitness when it did) ranged from -15% to 17%. Trends were similar in the forest (Figure 3), with significant differences among strains (F = 3.7, df = 10,17, p = 0.009), but no significant differences between killer and cured phenotypes or the interaction between phenotype and strain (phenotype: F = 0.29, df = 1,17, p = 0.60; interaction: F = 1.61, df = 10,17, p = 0.19). Similarly, naturally occurring killer and nonkiller S. paradoxus isolates, on average, had the same fitness in the forest (F = 0.16, df = 1,20.22, p = 0.70) (Figure 4). In the field assays, ddPCR reads decreased over time (t = 10.93, df = 168.75, p < 0.0001; Student’s t-test) (Figure 5), indicating a loss of cells; as a result, frequency changes in the forest should be interpreted as frequency changes during cell death, while fitness in the lab is reproductive fitness (Boynton et al., 2017; Vasi et al., 1994). Impacts of curing in the forest and the lab did not correlate with one another (Pearson’s correlation coefficient = 0.34, p = 0.51) (Figure 6).

**Figure 2.**
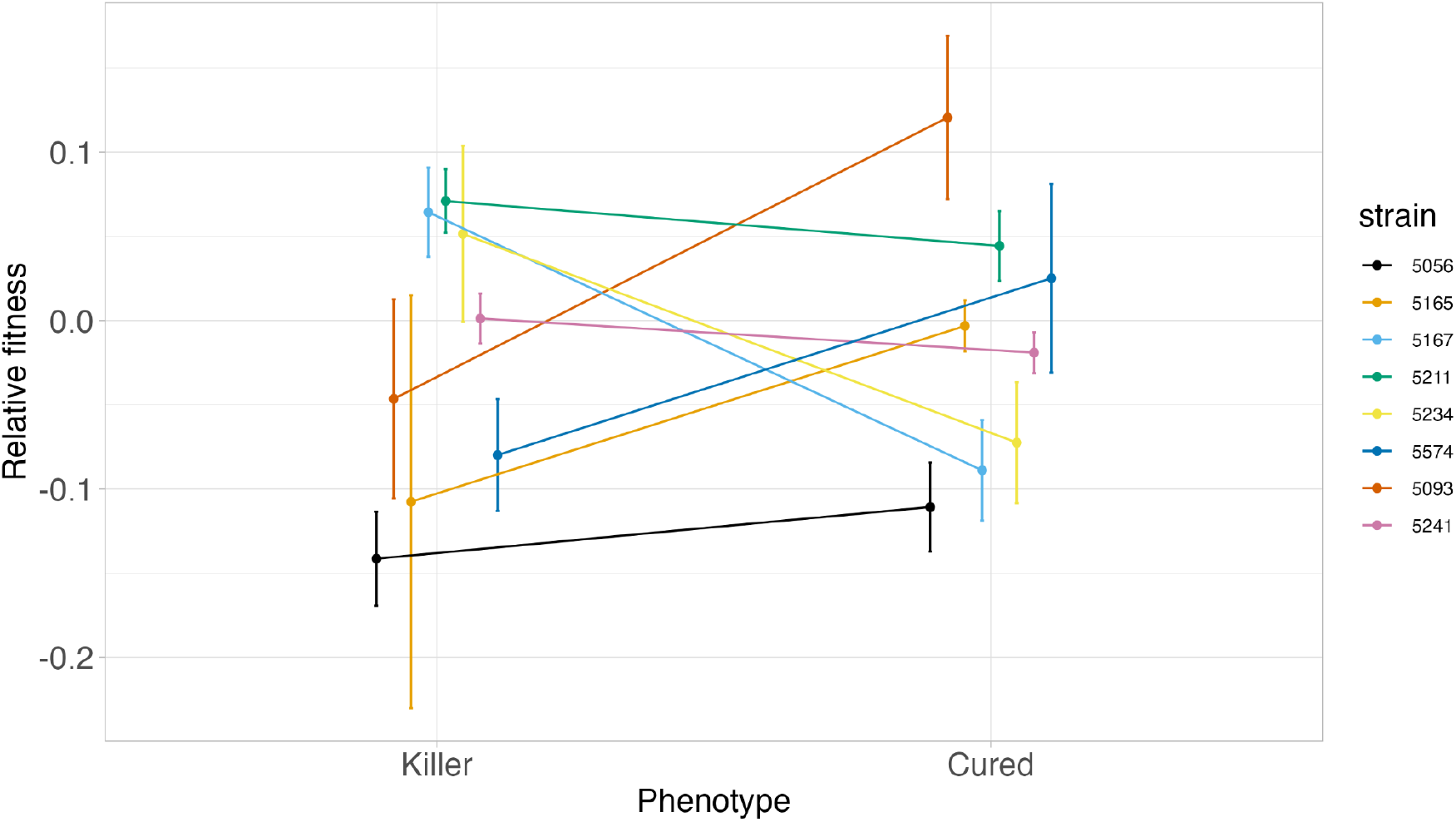
Virus curing had different impacts on the fitness of different S. paradoxus strains in the laboratory. Relative fitness of different tested strains relative to the reference strain is plotted on the y-axis, with colours denoting different strains. Each data point is the mean of up to six replicates for a tester strain, and the error bars show the standard error. Killer and cured versions of the same strain are joined by coloured lines.

**Figure 3.**
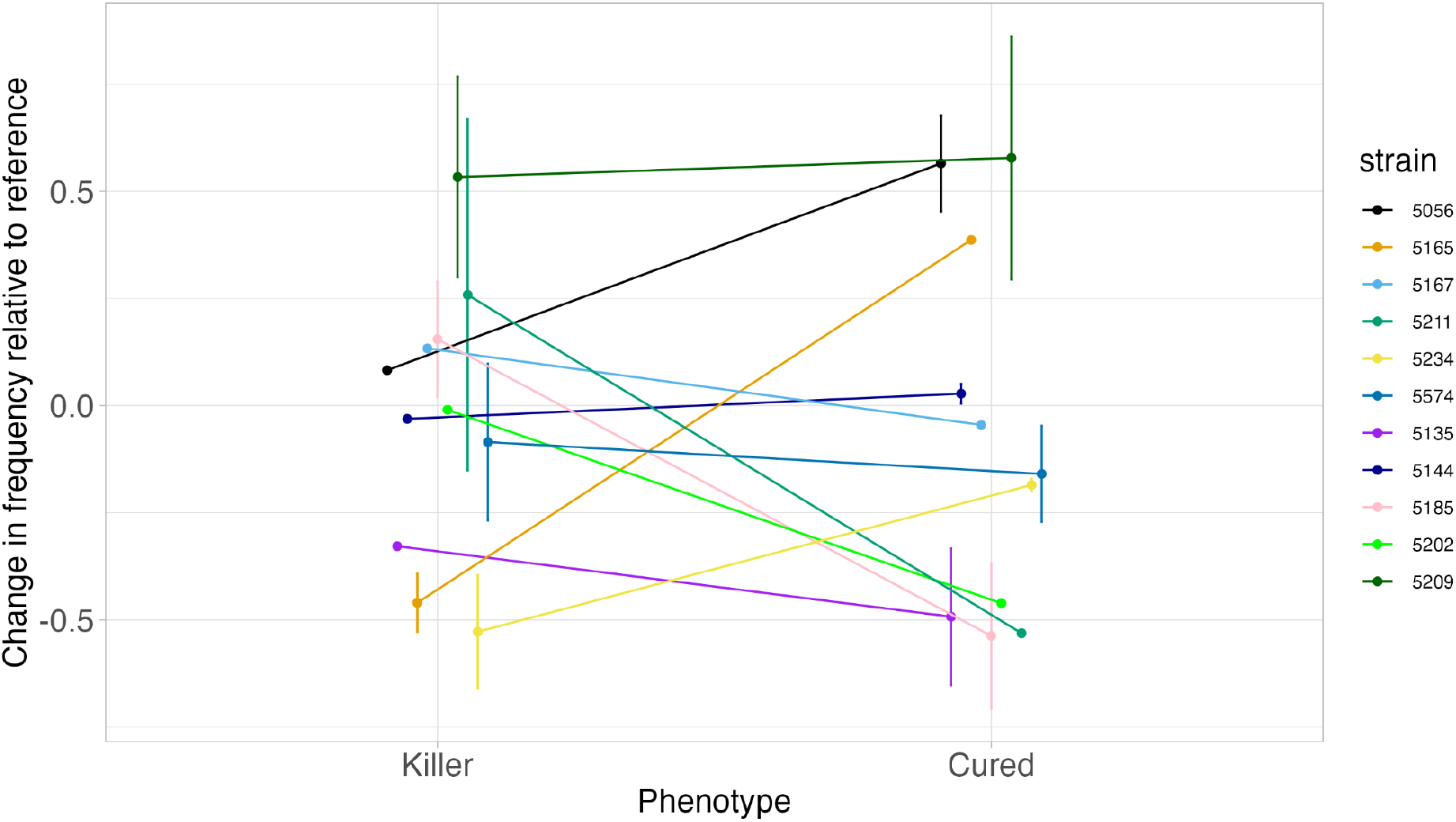
Strain identity determined the difference in fitness of S. paradoxus strains in the forest after curing. Relative fitness of the strains, as measured by change in frequency during the fitness assay relative to the reference strains, is plotted on the y-axis, and each data point is the mean of up to three replicates for each strain. Error bars indicate standard error. Strains are colour-coded, and killer and cured versions of the same strain are joined by coloured lines.

**Figure 4.**
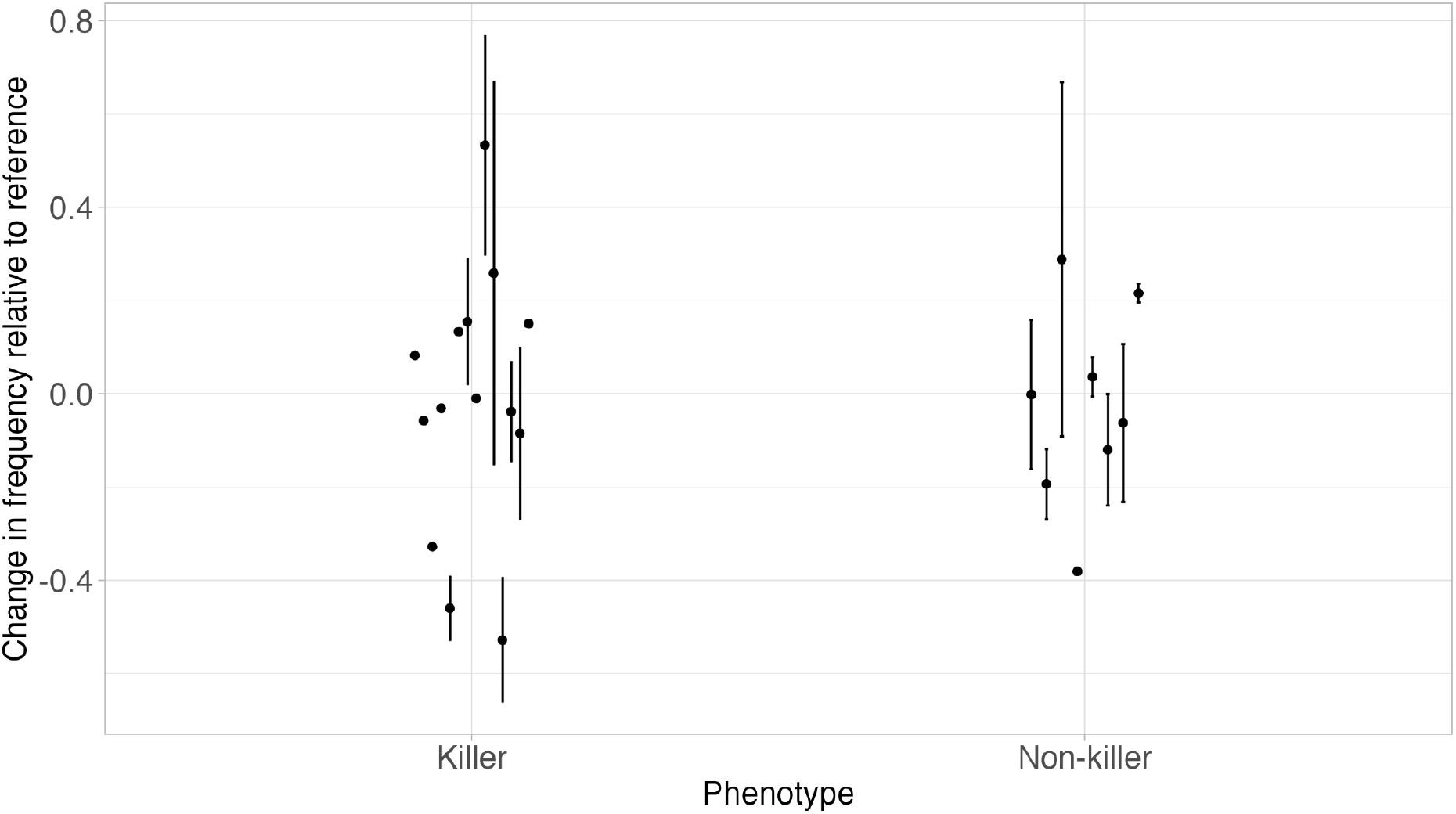
Killer and non-killer S. paradoxus strains do not show significantly different fitness in the wild. Relative fitness, as measured by the change in frequency relative to the reference during the fitness assay, is plotted on the y-axis. Each data point represents the mean relative fitness of up to three replicates and error bars represent the standard error.

**Figure 5.**
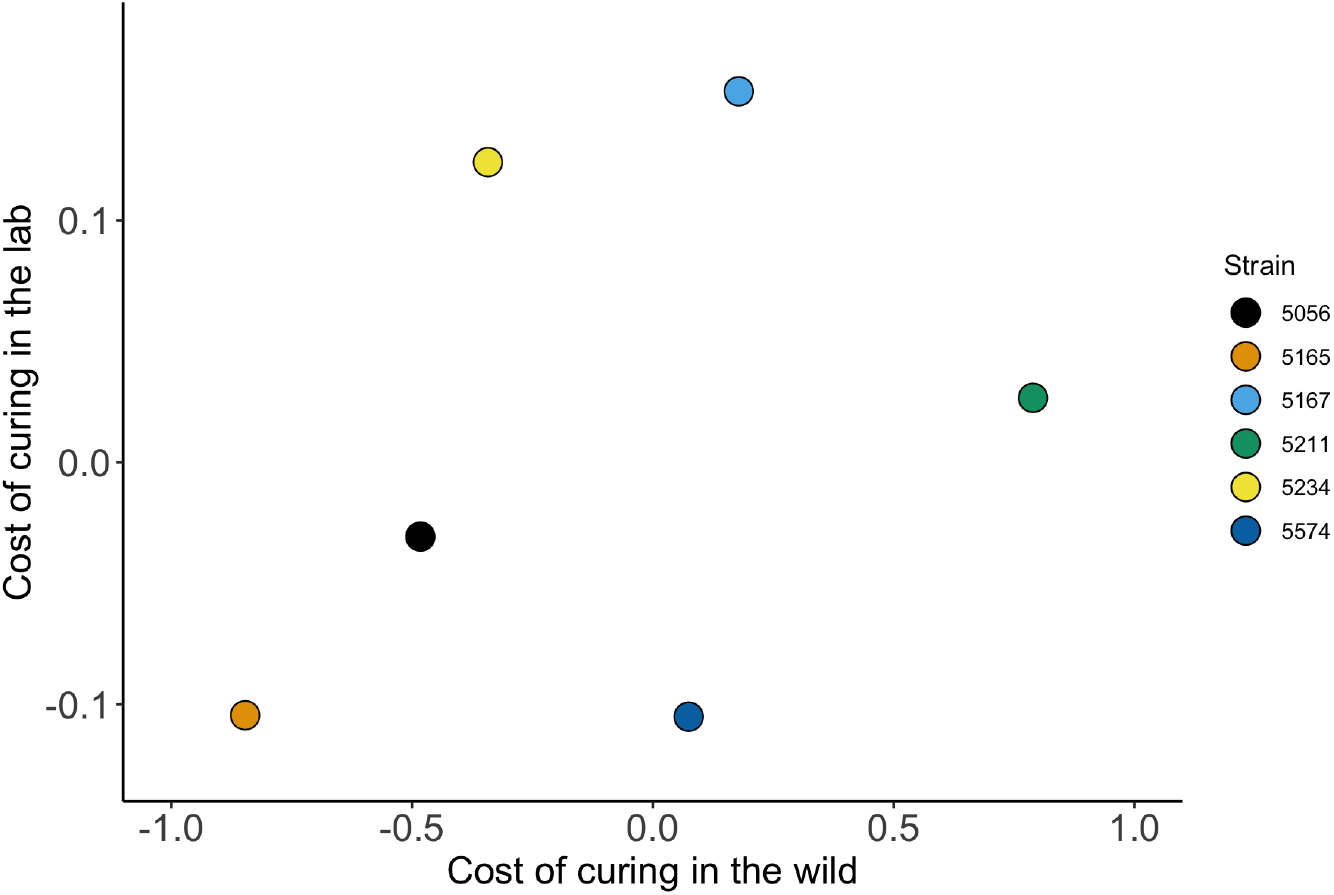
Change in host fitness upon curing (measured as the difference between fitness before and after the curing) in the laboratory and forest do not correlate. Only the six strains for which the tests were performed in both environments are included here; coloured dots indicate strain identity.

**Figure 6.**
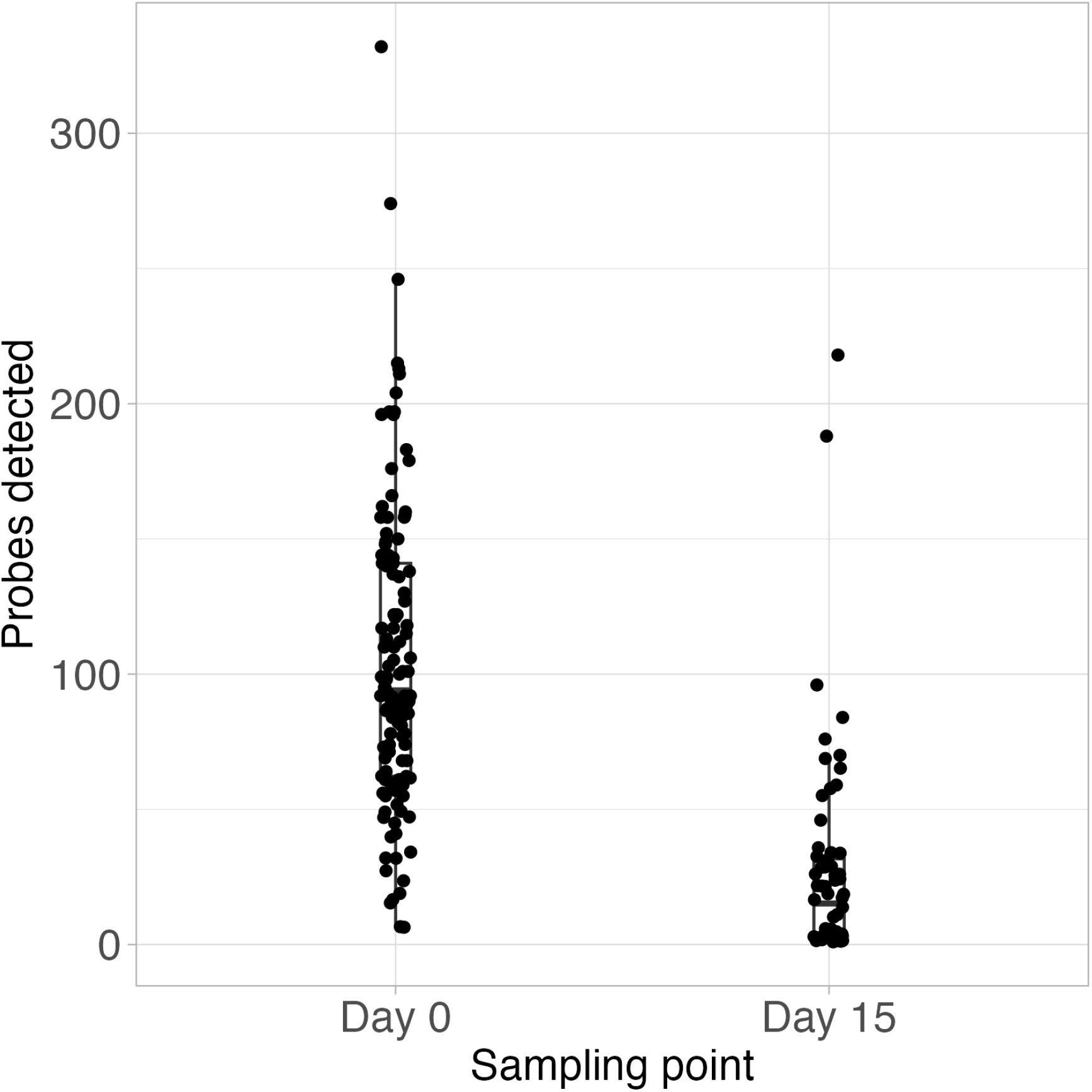
The number of detected probes significantly decreased during the fitness assay in the forest, from a mean of 104.9 at day 0 to a mean of 26.8 at day 15; t(168.75) = 10.93, p < 0.05, Student’s t-test. Probes that had concentration below 0.4 copies/ul were discarded from this analysis.

## Discussion

### Environmental conditions and host identity determine fitness consequences of virus removal

Killer virus impacts on host fitness vary depending on the host’s identity and the environment. In laboratory fitness assays, curing a wild *S. paradoxus* cell of its killer phenotype could have any possible fitness impact: viruses acted as parasites, commensals, and mutualists in different host backgrounds in the laboratory environment (Figure 2). In the field, while our methods could detect differences in overall strain fitness, we did not detect any fitness consequences of virus curing, suggesting that if fitness consequences are present, they are less impactful than natural fitness variation among *S. paradoxus* strains (Figure 3). Trends in fitness in the forest suggested a similar diversity of fitness impacts in the forest as in the lab. However, trends in fitness in the forest did not correlate with the effects of virus curing on fitness in the laboratory (Figure 5). Overall, our results suggest that researchers interested in the consequences of endosymbioses should carefully consider the genetic and environmental context of the endosymbiosis when designing tests.

Differences in strains’ responses to virus curing in the laboratory highlight diversity in evolution and coevolution among hosts and viruses. Killer viruses are likely to impose some fitness costs, including amino acid and other protein synthesis resources needed to produce secreted killer toxins (Pintar & Starmer, 2003). A history of host adaptation to viruses, potentially including compensatory adaptation to costs imposed by viruses, is a likely cause of observed fitness losses after curing in some *S. paradoxus* isolates (Pieczynska et al., 2017). Killer virus curing might also indirectly impact host growth by impacting helper virus copy numbers (Ball et al., 1984), although we did not investigate helper viruses directly in our experiment. We speculate that the observations made here reflect the diverse potential coevolutionary histories among interacting partners in the wild *S. paradoxus* cell.

Our results contrast with a previous observation of consistent wild *S. paradoxus* fitness loss after virus curing in laboratory conditions (Pieczynska et al., 2017). We interpret this contrast as another illustration of the varied nature of killer fitness effects: the previous observation included four wild *S. paradoxus* isolates from the United Kingdom, in contrast with our fourteen isolates from Germany; while both sets of strains are from the European *S. paradoxus* population, they represent different locations and slightly different evolutionary histories (Boynton et al., 2021), likely also having different coevolutionary histories with intracellular viruses. The few studies of fitnesses of cured lab-adapted killer *S. cerevisiae* show similar diversity in consequences of killing to our wild *S. paradoxus* in lab environments (Buskirk et al., 2020; Pieczynska et al., 2017), suggesting that coevolutionary history varies in length of time and number of adaptations across *Saccharomyces* populations.

Environmental dependence of fitness consequences is unsurprising in light of well-documented impacts of environments on microbial relative fitness. For example, genotype-by-environment interactions and local adaptation are frequent in *Saccharomyces* yeasts and many other organisms (Flynn et al., 2020; Kraemer & Boynton, 2017; Peltier et al., 2018). In our experiment, the contrast between the homogeneity of our well-mixed liquid laboratory cultures and the heterogeneity of our field chambers likely played a role in producing different patterns among environments (Figure 5): each field growth chamber replicate existed in a slightly different abiotic environment, which increased variation among replicates in our experiment. The phenotypes responsible for fitness diversity in both environments remain uncharacterized, and are targets of future research. Aside from toxin production, there are few documented phenotypes associated with intracellular yeast viruses, with the possible exception of changes in sporulation (Ravoitytė et al., 2020; Travers-Cook et al., 2022).

### Toxin production may further impact community interactions and fitness

Our experiments did not directly test the impacts of killer toxins on yeast competition in nature, which remain an open question. Our tests were designed to investigate the consequences of viral infection (and associated virus-sourced toxin production) in the *absence* of toxin-sensitive cells: no sensitive cells were present either in laboratory competitions or inside field chambers. While we did not collect any data on toxin impacts or activity, the effects of the killer toxin itself, especially in contact with or close proximity to sensitive cells, is likely to increase competitive fitness in certain environments. Toxin production has been hypothesised to be one of the main benefits of housing killer viruses, and to counteract hypothesised fitness costs of hosting killer viruses (Boynton, 2019; Pintar & Starmer, 2003). Toxin production can increase fitness in laboratory microcosms with sensitive cells, especially if killer virus-hosting cells are present in high frequencies and the abiotic environment is permissive to virus action (Buskirk et al., 2020; Greig & Travisano, 2008; Wloch-Salamon et al., 2008).

Evidence for toxin effectiveness in nature instead mostly comes from observational studies, which have produced conflicting results. We previously found no correlations between killer and toxin-resistant *S. paradoxus* in the same studied forest as the current study (Boynton et al., 2021), although others have found some correlations in other systems (Ganter & Starmer, 1992; Starmer et al., 1987). By isolating our killer and cured isolates from toxin-sensitive cells, we removed the impacts of toxins on competition from our tests so as to only test direct impacts of virus infection on host cells. Our hope is that studies like ours, which attempt to control for toxin activity, can provide a useful baseline for endosymbiont impacts on hosts on which phenotypes such as toxin production can be further studied.

### Microbial fitness can and should be investigated in natural field conditions

Our method of directly testing impacts of virus curing *in the field* opens up possibilities to answer several tantalising questions that we hope to answer going forward. Compared to traditional laboratory fitness assays, chamber experiments provide more realistic natural environments in which to explore fitness, but also can introduce environmental noise into experiments. Pairing field and laboratory assays resolves some of this tension. Because of the scale of our experiment, we saw a decrease in total ddPCR reads over experimental time in the forest (Figure 6), suggesting that the number of cells inoculated into chambers was higher than the carrying capacity of the forest environment. While this scenario did give detectable differences in fitness among *S. paradoxus* isolates, the relative fitnesses of *S. paradoxus* isolates, and of killer and cured *S. paradoxus*, are likely to be different in environments in which the number of inoculated cells is equal to (a likely scenario in nature) or below the environment’s carrying capacity. Similarly, we designed the experiment to exclude toxin-sensitive yeast cells, leaving the question open as to how effective killer toxins are for interference competition under natural conditions. Toxin effectiveness remains a key question in the yeast killer system (Boynton, 2019; Travers-Cook et al., 2023). Future applications of our chamber design are planned to include sensitive cells and to monitor microbial community changes outside of chambers. To our knowledge, neither the community-wide impacts of killer toxin production nor the distance at which toxins can diffuse and maintain effectiveness have been studied. We intend for the current experiment to open doors to future investigations of the ecological costs and benefits to hosting intracellular endosymbionts, both in the killer yeast system and with other microbial hosts.

## Conclusion

Research on the consequences of endosymbiosis of microbial hosts *in natural contexts* is in its infancy. The tractability of the *Saccharomyces* killer system has made it an ideal model system in which to study toxin-related molecular interactions and evolution in controlled laboratory conditions (Martinac et al., 1990; McBride et al., 2008; Pieczynska et al., 2016), but yeast researchers don’t know much about the consequences of the endosymbiosis itself over millions of years of yeast evolution (Liti et al., 2006; Shen et al., 2018). Here we used field-based chambers to compare endosymbionts’ influence on fitness in nature with laboratory measurements.

Maintenance of killer virus-like endosymbioses in nature is likely due to a combination of factors that we did test, including costs of carrying endosymbionts and coevolutionary adaptation to endosymbiont presence, combined with factors that we did not test, including direct impacts of host toxins on the microbial community and host-symbiont interactions that do not directly change fitness relative to resistant controls (*e*.*g*., symbionts behaving as selfish toxin-antitoxin systems) (Kast et al., 2015; Travers-Cook et al., 2023). Going forward, our field chamber system will allow for the breakdown and testing of each of these mechanisms, assembling a complete picture of endosymbiosis maintenance in yeast and similar systems. Importantly, chambers like ours can be used for similar experiments with any microbial host organism, and we hope future researchers will use them to ask similar questions with other model and non-model microbes. *Saccharomyces* yeasts are an ideal model system for understanding endosymbiosis, and our results highlight that microbial hosts can have as complex interactions with intracellular endosymbionts as any large multicellular organism in its native environment.

## Data availability

All data for Figures 2, 3, 4, and 6 are included in the supplementary information. Data for Figure 5 is based on data for Figures 2 and 3.

## Acknowledgements

The authors are grateful for helpful discussions and laboratory support from Doreen Landermann, Benjamin Claassen, Sarah Frohriep, and Ezgi Özkurt, and helpful discussions from Dominkia Wloch-Salamon, Daniel Unterweger, and Paul Rowley. We thank Duncan Greig and Dominika Wloch-Salamon for access to tester *Saccharomyces cerevisiae* strains. This research was funded by a Max Planck Fellowship to EHS.

## Author contributions

Conceptualization: RU, PJB; Data curation: RU; Formal analysis: RU; Funding acquisition: EHS; Investigation: RU; Methodology: RU, OEK, PJB; Project administration: PJB; Resources: EHS; Supervision: EHS, PJB; Validation: PJB; Visualization: RU, PJB; Writing–original draft: RU, PJB; Writing–review & editing: RU, OEK, EHS, PJB

---

**Supplementary table 1.**

| Strain (genotype) | Phenotype | Used in lab/wild fitness assay |
| --- | --- | --- |
| 5241 (*hxt3-1*) | Killer | lab |
| 5056 (*hxt3-1*) | Killer | lab & wild |
| 5202 (*hxt3-1*) | Killer | wild |
| 5209 (*hxt3-1*) | Killer | wild |
| 5135 (*hxt3-1*) | Killer | wild |
| 5144 (*hxt3-1*) | Killer | wild |
| 5185 (*hxt3-1*) | Killer | wild |
| 5587 (*hxt3-2*) | Killer | wild |
| 5211 (*hxt3-2*) | Killer | lab & wild |
| 5167 (*hxt3-2*) | Killer | lab & wild |
| 5165 (*hxt3-2*) | Killer | lab & wild |
| 5234 (*hxt3-2*) | Killer | lab & wild |
| 5093 (*hxt3-2*) | Killer | lab |
| 5574 (*hxt3-2*) | Killer | lab & wild |
| 6622 (*hxt3-1*) | Cured | lab |
| 6804 (*hxt3-1*) | Cured | lab & wild |
| 6805 (*hxt3-1*) | Cured | wild |
| 6806 (*hxt3-1*) | Cured | wild |
| 6808 (*hxt3-1*) | Cured | wild |
| 6811 (*hxt3-1*) | Cured | wild |
| 6812 (*hxt3-1*) | Cured | wild |
| **6813 (hxt3-2)** | **Cured** | **wild** |
| **6619 (hxt3-2)** | **Cured** | **lab & wild** |
| **6620 (hxt3-2)** | **Cured** | **lab & wild** |
| **6621 (hxt3-2)** | **Cured** | **lab & wild** |
| **6807 (hxt3-2)** | **Cured** | **lab & wild** |
| **6809 (hxt3-2)** | **Cured** | **lab** |
| **6810 (hxt3-2)** | **Cured** | **lab & wild** |
| **5119 (hxt3-1)** | **Non-killer** | **wild** |
| **5203 (hxt3-1)** | **Non-killer** | **wild** |
| **5236 (hxt3-1)** | **Non-killer** | **wild** |
| **5222 (hxt3-1)** | **Non-killer** | **wild** |
| **5221 (hxt3-1)** | **Non-killer** | **wild** |
| **5212 (hxt3-2)** | **Non-killer** | **wild** |
| **5204 (hxt3-2)** | **Non-killer** | **wild** |
| **5087 (hxt3-2)** | **Non-killer** | **wild** |
| **5164 (hxt3-2)** | **Non-killer** | **wild** |
| **5227 (hxt3-2)** | **Non-killer** | **wild** |
| **5149 (hxt3-2)** | **Non-killer** | **wild (reference for batch 1)** |
| **5235 (hxt3-1)** | **Non-killer** | **wild (reference for batch 2)** |
| **5149t (hxt3-2)** | **Non-killer** | **lab (reference; transformed with kan-resistance)** |

## Notes

### Competing Interest Statement

The authors have declared no competing interest.

## Citations

Ball, S. G., Tirtiaux, C., & Wickner, R. B. (1984). Genetic control of L-A and L-(BC) dsRNA copy number in killer systems of Saccharomyces cerevisiae. Genetics, 107(2), 199–217. 10.1093/genetics/107.2.199

Bates, D., Mächler, M., Bolker, B., & Walker, S. (2015). Fitting linear mixed-effects models using lme4. Journal of Statistical Software, 67, 1–48. 10.18637/jss.v067.i01

Boynton, P. J. (2019). The ecology of killer yeasts: Interference competition in natural habitats. Yeast, 36(8), 473–485. 10.1002/yea.3398

Boynton, P. J., & Greig, D. (2014). The ecology and evolution of non-domesticated Saccharomyces species. Yeast, 31(12), 449–462. 10.1002/yea.3040

Boynton, P. J., Stelkens, R., Kowallik, V., & Greig, D. (2017). Measuring microbial fitness in a field reciprocal transplant experiment. Molecular Ecology Resources, 17(3), 370–380. 10.1111/1755-0998.12562

Boynton, P. J., Wloch-Salamon, D., Landermann, D., & Stukenbrock, E. H. (2021). Forest Saccharomyces paradoxus are robust to seasonal biotic and abiotic changes. Ecology and Evolution, 11(11), 6604–6619. 10.1002/ece3.7515

Bright, M., & Bulgheresi, S. (2010). A complex journey: Transmission of microbial symbionts. Nature Reviews Microbiology, 8(3), 218–230. 10.1038/nrmicro2262

Buskirk, S. W., Rokes, A. B., & Lang, G. I. (2020). Adaptive evolution of nontransitive fitness in yeast. eLife, 9, e62238. 10.7554/eLife.62238

Castledine, M., & Buckling, A. (2024). Critically evaluating the relative importance of phage in shaping microbial community composition. Trends in Microbiology. 10.1016/j.tim.2024.02.014

Crabtree, A. M., Taggart, N. T., Lee, M. D., Boyer, J. M., & Rowley, P. A. (2023). The prevalence of killer yeasts and double-stranded RNAs in the budding yeast Saccharomyces cerevisiae. FEMS Yeast Research, 23, foad046. 10.1093/femsyr/foad046

del Arco, A., Fischer, M. G., & Becks, L. (2024). Evolution of exploitation and replication of giant viruses and virophages. Virus Evolution, 10(1), veae021. 10.1093/ve/veae021

El-Khatib, S., Lambert, M. G., Reed, M. N., Goncalves, M. B., & Boynton, P. J. (2023). Leaf decomposing fungi influence Saccharomyces paradoxus growth across carbon environments. microPublication Biology. 10.17912/micropub.biology.000739

Fink, G. R., & Styles, C. A. (1972). Curing of a killer factor in Saccharomyces cerevisiae. Proceedings of the National Academy of Sciences, 69(10), 2846–2849. 10.1073/pnas.69.10.2846

Flynn, J. M., Rossouw, A., Cote-Hammarlof, P., Fragata, I., Mavor, D., Hollins, C., III, Bank, C., & Bolon, D. N. (2020). Comprehensive fitness maps of Hsp90 show widespread environmental dependence. eLife, 9, e53810. 10.7554/eLife.53810

Fry, A. J., Palmer, M. R., & Rand, D. M. (2004). Variable fitness effects of Wolbachia infection in Drosophila melanogaster. Heredity, 93(4), 379–389. 10.1038/sj.hdy.6800514

Gabaldón, T. (2021). Origin and early evolution of the eukaryotic cell. Annual Review of Microbiology, 75, 631–647. 10.1146/annurev-micro-090817-062213

Ganter, P. F., & Starmer, W. T. (1992). Killer factor as a mechanism of interference competition in yeasts associated with cacti. Ecology, 73(1), 54–67. 10.2307/1938720

Gietz, D. R., & Woods, R. A. (2002). Transformation of yeast by lithium acetate/single-stranded carrier DNA/polyethylene glycol method. In C. Guthrie & G. R. Fink (Eds.), Methods in Enzymology (Vol. 350, pp. 87–96). Academic Press. 10.1016/S0076-6879(02)50957-5

Greig, D., & Travisano, M. (2008). Density-dependent effects on allelopathic interactions in yeast. Evolution, 62(3), 521–527. 10.1111/j.1558-5646.2007.00292.x

Hassani, M. A., Özkurt, E., Seybold, H., Dagan, T., & Stukenbrock, E. H. (2019). Interactions and coadaptation in plant metaorganisms. Annual Review of Phytopathology, 57(Volume 57, 2019), 483–503. 10.1146/annurev-phyto-082718-100008

Johnson, N. c. (1998). Responses of Salsola kali and Panicum virgatum to mycorrhizal fungi, phosphorus and soil organic matter: Implications for reclamation. Journal of Applied Ecology, 35(1), 86–94. 10.1046/j.1365-2664.1998.00277.x

Kast, A., Voges, R., Schroth, M., Schaffrath, R., Klassen, R., & Meinhardt, F. (2015). Autoselection of cytoplasmic yeast virus-like elements encoding toxin/antitoxin systems involves a nuclear barrier for immunity gene expression. PLOS Genetics, 11(5), e1005005. 10.1371/journal.pgen.1005005

Kraemer, S. A., & Boynton, P. J. (2017). Evidence for microbial local adaptation in nature. Molecular Ecology, 26(7), 1860–1876. 10.1111/mec.13958

Kuznetsova, A., Brockhoff, P. B., & Christensen, R. H. B. (2017). lmerTest Package: Tests in linear mixed effects models. Journal of Statistical Software, 82, 1–26. 10.18637/jss.v082.i13

Lenski, R. E., Rose, M. R., Simpson, S. C., & Tadler, S. C. (1991). Long-term experimental evolution in Escherichia coli. I. Adaptation and divergence during 2,000 generations. The American Naturalist, 138(6), 1315–1341. 10.1086/285289

Liti, G., Barton, D. B. H., & Louis, E. J. (2006). Sequence diversity, reproductive isolation and species concepts in Saccharomyces. Genetics, 174(2), 839–850. 10.1534/genetics.106.062166

Lloyd, G. S., & Thomas, C. M. (2023). Microbial primer: The logic of bacterial plasmids. Microbiology, 169(7), 001336. 10.1099/mic.0.001336

Lozupone, C. A., Stombaugh, J. I., Gordon, J. I., Jansson, J. K., & Knight, R. (2012). Diversity, stability and resilience of the human gut microbiota. Nature, 489(7415), 220–230. 10.1038/nature11550

Martinac, B., Zhu, H., Kubalski, A., Zhou, X. L., Culbertson, M., Bussey, H., & Kung, C. (1990). Yeast K1 killer toxin forms ion channels in sensitive yeast spheroplasts and in artificial liposomes. Proceedings of the National Academy of Sciences, 87(16), 6228–6232. 10.1073/pnas.87.16.6228

McBride, R., Greig, D., & Travisano, M. (2008). Fungal viral mutualism moderated by ploidy. Evolution, 62(9), 2372–2380. 10.1111/j.1558-5646.2008.00443.x

Obeng, N., Pratama, A. A., & van Elsas, J. D. (2016). The significance of mutualistic phages for bacterial ecology and evolution. Trends in Microbiology, 24(6), P440–449. 10.1016/j.tim.2015.12.009

Peltier, E., Sharma, V., Martí Raga, M., Roncoroni, M., Bernard, M., Jiranek, V., Gibon, Y., & Marullo, P. (2018). Dissection of the molecular bases of genotype x environment interactions: A study of phenotypic plasticity of Saccharomyces cerevisiae in grape juices. BMC Genomics, 19(1), 772. 10.1186/s12864-018-5145-4

Pieczynska, M. D., Korona, R., & De Visser, J. A. G. M. (2017). Experimental tests of host–virus coevolution in natural killer yeast strains. Journal of Evolutionary Biology, 30(4), 773–781. 10.1111/jeb.13044

Pieczynska, M. D., Wloch-Salamon, D., Korona, R., & de Visser, J. A. G. M. (2016). Rapid multiple-level coevolution in experimental populations of yeast killer and nonkiller strains. Evolution, 70(6), 1342–1353. 10.1111/evo.12945

Pinheiro, J., Bates, D., DebRoy, S., Sarkar, D., & R Core Team. (2018). nlme: Linear and Nonlinear Mixed Effects Models [Computer software]. https://CRAN.R-project.org/package=nlme

Pintar, J., & Starmer, W. T. (2003). The costs and benefits of killer toxin production by the yeast Pichia kluyveri. Antonie van Leeuwenhoek, 83(1), 89–97. 10.1023/A:0000000089097

R Core Team. (2017). R: A language and environment for statistical computing (Version 3.4.3) [Computer software]. R Foundation for Statistical Computing. https://www.R-project.org/

Ravoitytė, B., Lukša, J., Yurchenko, V., Serva, S., & Servienė, E. (2020). Saccharomyces paradoxus transcriptional alterations in cells of distinct phenotype and viral dsRNA content. Microorganisms, 8(12), Article 12. 10.3390/microorganisms8121902

Rowley, P. A. (2017). The frenemies within: Viruses, retrotransposons and plasmids that naturally infect Saccharomyces yeasts. Yeast, 34(7), 279–292. 10.1002/yea.3234

San Millan, A., & MacLean, R. C. (2017). Fitness costs of plasmids: A limit to plasmid transmission. Microbiology Spectrum, 5(5), 10.1128/microbiolspec.mtbp-0016-2017. https://doi.org/10.1128/microbiolspec.mtbp-0016-2017

Schmitt, M. J., & Breinig, F. (2006). Yeast viral killer toxins: Lethality and self-protection. Nature Reviews Microbiology, 4(3), 212–221. 10.1038/nrmicro1347

Shapiro, J. W., & Turner, P. E. (2018). Evolution of mutualism from parasitism in experimental virus populations. Evolution, 72(3), 707–712. 10.1111/evo.13440

Shen, X.-X., Opulente, D. A., Kominek, J., Zhou, X., Steenwyk, J. L., Buh, K. V., Haase, M. A. B., Wisecaver, J. H., Wang, M., Doering, D. T., Boudouris, J. T., Schneider, R. M., Langdon, Q. K., Ohkuma, M., Endoh, R., Takashima, M., Manabe, R., Čadež, N., Libkind, D., … Rokas, A. (2018). Tempo and mode of genome evolution in the budding yeast subphylum. Cell, 175(6), 1533-1545.e20. 10.1016/j.cell.2018.10.023

Starmer, W. T., Ganter, P. F., Aberdeen, V., Lachance, M.-A., & Phaff, H. J. (1987). The ecological role of killer yeasts in natural communities of yeasts. Canadian Journal of Microbiology, 33(9), 783–796. 10.1139/m87-134

Travers-Cook, T. J., Jokela, J., & Buser, C. C. (2023). The evolutionary ecology of fungal killer phenotypes. Proceedings of the Royal Society B: Biological Sciences, 290(2005), 20231108. 10.1098/rspb.2023.1108

Travers-Cook, T. J., Skirgaila, C., Martin, O. Y., & Buser, C. C. (2022). Infection by dsRNA viruses is associated with enhanced sporulation efficiency in Saccharomyces cerevisiae. Ecology and Evolution, 12(1), e8558. 10.1002/ece3.8558

Vasi, F., Travisano, M., & Lenski, R. E. (1994). Long-term experimental evolution in Escherichia coli. II. Changes in life-history traits during adaptation to a seasonal environment. The American Naturalist, 144(3), 432–456. 10.1086/285685

Wang, Q.-M., Liu, W.-Q., Liti, G., Wang, S.-A., & Bai, F.-Y. (2012). Surprisingly diverged populations of Saccharomyces cerevisiae in natural environments remote from human activity. Molecular Ecology, 21(22), 5404–5417. 10.1111/j.1365-294X.2012.05732.x

Wickham, H. (2016). ggplot2: Elegant Graphics for Data Analysis (2nd ed.). Springer-Verlag. https://ggplot2.tidyverse.org

Wickner, R. B. (1996). Double-stranded RNA viruses of Saccharomyces cerevisiae. Microbiological Reviews, 60(1), 250–265. 10.1128/mr.60.1.250-265.1996

Wloch-Salamon, D. M., Gerla, D., Hoekstra, R. F., & de Visser, J. A. G. M. (2008). Effect of dispersal and nutrient availability on the competitive ability of toxin-producing yeast. Proceedings of the Royal Society B: Biological Sciences, 275(1634), 535–541. 10.1098/rspb.2007.1461

